# Reticulate Evolutionary History of a Western Palaearctic Bat Complex Explained by Multiple mtDNA Introgressions in Secondary Contacts

**DOI:** 10.1101/348235

**Authors:** Emrah çoraman, Christian Dietz, Elisabeth Hempel, Astghik Gazaryan, Eran Levin, Primož Presetnik, Maja Zagmajster, Frieder Mayer

## Abstract

**Aim:** There is increasing evidence showing that species within various taxonomic groups have reticulate evolutionary histories with several cases of introgression events. Investigating the phylogeography of species complexes can provide insight about the introgressions, when and where these hybridizations occurred. In this study, we investigate the biogeography of a widely distributed Western Palaearctic bat species complex, namely *Myotis nattereri* sensu lato. This complex exhibits high genetic diversity and in its western distribution range is composed of deeply diverged genetical lineages. However, little is known about the genetic structure of the eastern populations. We also infer the conservation and taxonomical implications of the identified genetic divergences.

**Location:** Western Palaearctic

**Methods:** We analyzed 175 specimens collected from 67 locations and sequenced one mitochondrial and four nuclear DNA markers, and combined these with the available GenBank sequences. We used haplotype networks, PCA, t-SNE, and Bayesian clustering algorithms to investigate the population structure and Bayesian trees to infer the phylogenetic relationship of the lineages.

**Main conclusions:** We identified deeply divergent genetical lineages. In some cases, nuclear and mitochondrial markers were discordant, which we interpret are caused by hybridization between lineages. We identified three such introgression events. Our findings suggest that the M. *nattereri* complex has a reticulate evolutionary history with multiple cases of hybridizations between some of the identified lineages. We also suggest a revision in the taxonomy of this species group, with two possible new taxa: *M. hoveli* and *M. tschuliensis.*

## 1 Introduction

Hybridization events are critical milestones in the evolutionary history of species. Whenever individuals from two distinct taxa successfully produce offspring, there occurs a chance that their genes could be introduced into the gene pool of the other’s. If both parental populations are large and the number of hybrids are few, it is likely that the genetic contribution of such gene flow will be minute and probably will quickly diminish because of genetic drift. However, if one of the populations is small or the introduced genes have a selective advantage, then it is more likely that the introduced genes can spread in their new host population. Nowadays, there is increasing evidence, showing that introgressions occurred in diverse taxonomic groups. For example, birds (Rheindt and Edwards 2011), cats (Li et al. 2016), freshwater fishes (Wallis et al. 2017), and humans (Prüfer et al. 2014) exhibit reticulate evolutionary histories, with multiple cases of introgression events. These findings suggest that hybridization between distinct taxa occurred more than occasionally, playing a prominent role in shaping the extant genetic structure of various species groups (Toews and Brelsford 2012).

Investigating the biogeographical history of species complexes can help us to understand the underlying mechanism of introgressions. Hybridizations are likely to happen when species colonize new areas (Currat et al. 2008); when a population extends its range, it can get into contact with another local taxa, such as a closely related species or another conspecific lineage. These contacts provide a chance for hybridizations and consequently, introgressions. First invaders would be presumably males and more likely to find a mate from the local population. In such a case, first hybrids would have the mitochondrial DNA (mtDNA) of the local population, and their nuclear genome from both of their parents. If, in the successive generations, these hybrids would backcross with the newcomers, their nuclear genome would homogenize with the invaders. In such a scenario, even low levels of gene flow can result in introgression of alleles, and even in the lack of selective advantage (Currat et al. 2008; Petit and Excoffier 2009). In case of male dominated invasions, uniparentally inherited organelle DNA would even more quickly introgress, as the local populations would be their only source. In that sense, discordance between organelle and nuclear genomes would be an indication of possible introgression events (Currat et al. 2008; Toews and Brelsford 2012).

In this study, we investigate the phylogeographical patterns of a wide spread morphologically cryptic bat species complex by using mitochondrial and nuclear markers. Our study taxa, *Myotis nattereri* sensu lato, are distributed in Northwest Africa, Europe, and parts of the Middle East (Figure 1). In the western ranges, the group is composed of spatially structured genetic lineages, some of which are proposed as distinct species: *M. escaleari* in the Iberian peninsula (Ibáñez et al. 2006); *M*. species A in the Italian peninsula and around the Pyrenees (Salicini et al. 2011, 2013); M. species B in the Northwest Africa (Salicini et al. 2011, 2013); *M*. species C in Corsica (Puechmaille et al. 2012); and the rest of Europe is represented by the nominal form, *M. nattereri.* Relatively little is known for the eastern ranges. In parts of the Caucasus, the nominal form is sympatric with a larger form, *M. schaubi.* This larger form is clearly distinct both in morphology and in genetics (Ruedi and Mayer 2001; Salicini et al. 2011). Surprisingly, in mtDNA, the *M. nattereri* populations from southern Anatolia cluster with this species (Çoraman et al. 2013). Here, by adding samples mostly from the eastern distribution ranges, we investigate the phylogeographical patterns of the whole species group and infer its evolutionary history.

Our results show that the *M. nattereri* species group has a complex and reticulate evolutionary history involving multiple cases of hybridization between divergent genetic lineages. We identified unknown cryptic diversity in the eastern ranges. All of the identified genetic lineages evolved in distinct glacial refugia and later on, expanded their ranges during the interglacial periods. In some of these expansions, neighboring lineages got into contact and hybridized with each other. We identified three major hybridization events, which ended up with mitochondrial introgressions. In these cases, the expanding lineages got the mtDNA from the local populations, some of which got extinct but their mtDNA survived.

## 2 Material and Methods

### 2.1 Genetic Data Sampling

We analyzed 175 samples collected from 67 locations and added 158 samples from 92 locations gathered from GenBank (Figure 1 a and Table S1.1). DNA isolation and the sequencing of partial mitochondrial gene NADH dehydrogenase subunit 1 (ND1) was done as described in Dietz et al. (2016). For mtDNA analysis, we generated 166 sequences (ranging between 403 to 650 bp long) and analyzed them with 158 sequences gathered from GenBank (Table S1.2). For nuDNA analysis, we sequenced nuclear introns following Salicini et al. (2011, 2018), whom previously used these markers for this species group (Table 1). We selected four markers, which were informative to identify the previous described lineages. Sequences were edited and aligned in CodonCode Aligner (CodonCode Corporation, MA, USA) and nuclear sequences were phased by using the PHASE algorithm as implemented in DNASP version 5.10 (Librado and Rozas 2009). Estimates of genetic divergence among lineages were conducted in MEGA7 (Kumar et al. 2016) and sequence polymorphism estimates in DNASP. We generated haplotype networks using the HaploNet function as implemented in the R package Rpegas v 0.10 (Paradis 2010). All the generated genetic data is deposited in GenBank (Table S1.1).

### 2.2 Phylogenetic Reconstructions

For the mtDNA analysis, we reconstructed a Bayesian phylogenetic tree using BEAST v1.8.3 (Drummond et al. 2012). We used 271 sequences (Table S1.1) and trimmed them to 499 bp long to have equal lengths. We ran three independent 25 million chains with UPGMA starting trees. Yule speciation prior was specified and HKY+G was selected as the substitution model, based the Bayesian Information Criterion as implemented in jModelTest v2.1.10 (Darriba et al. 2012). We checked the convergence of the runs and the effective sample size (ESS) of the estimated parameters in Tracer v1.6.0 (http://tree.bio.ed.ac.uk/software/tracer). In all runs, all the parameters converged and they had ESS values higher than 200. The resulting tree and log files of repeated runs were combined in LogCombiner v1.8.0 (http://beast.bio.ed.ac.uk/logcombiner) with a 10% burnin. The trees were summarized using the maximum clade credibility topology in TreeAnnotator and the final tree was rooted with *M. bechsteinii* and *M. emarginatus*. In order to identify the lineage identity of the samples which had shorter ND1 sequences, we also constructed a neighbor-joining tree with 500 replicates and using p-distances as implemented in MEGA7.

For the species tree reconstructions, we used two different approaches. In the first approach we used *BEAST (Heled and Drummond 2009) as implemented in the BEAST v1.8.3 (Drummond et al. 2012). Here we grouped the samples based on their mtDNA clade identities, but also introduced two new groups, populations which had discordant results in the PCA (see Results section 3.2): the *M. nattereri* lineage individuals, which clustered in the Central lineage; and the Eastern lineage individuals from southeastern Anatolia and Israel, which clustered together with the Central and the *M. nattereri* lineages. Few individuals representing each group were selected and the phased intron sequences were analyzed. We used Yule model for the tree prior and HKY as the substitution model. Three independent runs of 50 million chains were computed and post processed similar to the mtDNA BEAST runs.

In the second approach, we used the Bayesian Phylogenetics and Phylogeography (BPP) program in the unguided species delimitation mode (A11) as described in Yang (2015). The BPP program utilizes a Bayesian Markov chain Monte Carlo (MCMC) algorithm for analyzing DNA sequence alignments under the multispecies coalescent model, and in addition to species tree estimation can also infer species delimitation (Rannala and Yang 2003; Takahata et al. 1995; Yang 2002). We used the ‘algorithm 0’ for the RJalgorithm, and selected ‘uniform rooted trees’ for the species model prior with using the default parameters for Gamma, Gamma-Dirichlet, and MCMC variables. We repeated the runs three times and their results were consistent.

### 2.3 Population Genetic Structure

We used two approaches to investigate the population structure. In the first one, we ran principal component analyses (PCA) using the R package adegenet v 2.1.1 (Jombart 2008). We used samples which had data for all nuclear markers. We ran the analysis twice, both for the haplotype and for the concatenated variable sites.

As a second approach, we used t-distributed stochastic neighbor embedding (t-SNE) (van der Maaten and Hinton 2008) to qualitatively evaluate population structure. The t-SNE method is a dimension reduction and visualization technique for high-dimensional data, which have been also used for genetic data (Li et al. 2017). For this analysis, we used the R package Rtsne v 0.13 (https://github.com/jkrijthe/Rtsne). We converted the concatenated SNP data to the euclidean genetic distances, and set the perplexity to 10 and the maximum number of iterations to 5000.

## 3 Results

### 3.1 Mitochondrial Phylogeny

Bayesian reconstruction of the ND1 sequences revealed a deeply diverged phylogenetic tree. Its topography was similar to those previously reported (Salicini et al. 2011, 2013; Figure 1 b). First, the tree splits into two main branches, both of which further split into two, forming the four major lineage groups.

#### *M. nattereri* lineage

Among the identified lineages, the *M. nattereri* lineage has the widest distribution. Its range spreads from Ireland in the west to Ukraine in the east, and from the Mediterranean Sea in the south to the Baltic Sea in the north. Despite its wide distribution range, the *M. nattereri* lineage has relatively shallow genetic diversity (Table 2). The distribution of the haplotypes and their network structure suggest that its extant populations survived the last glacial period in three regions: i) western Balkans; ii) Greece; and iii) the western Anatolia (Figure S2.1 a). Based on the geographical distribution of the haplotypes, we identified two expansions out of the Balkans: one to the west and one to the east. Samples from Ireland, southern France, and Balkans shared the same haplotype, pointing to their very recent common ancestry.

#### Central lineage

The second major lineage is sister to the *M. nattereri* lineage, and composed of two distinct clades: i) *M.* sp A in northern Italy, Slovenia, and the Pyrenees; and ii) another one in Southern Italy (will be referred as the Southern Italy lineage). These clades differed by approximately 7% (Table 3) and had a contact zone in the central Apennine Mountains. The *M.* sp A lineage exhibit spatial structuring (Figure S2.1 b); populations around the Pyrenees were separated from the populations in the southern Alps, which suggested an earlier expansion into the Pyrenees.

#### Western lineage

The third major lineage is also composed of two clades: i) *M. escalerai* from the Iberian peninsula; and ii) *M*. sp B from Northern Africa. These clades differed around 8%

#### Eastern lineage

The fourth lineage was composed of the populations from the Caucasus, Crimea, and the coastal zones of the eastern Mediterranean. This lineage is highly structured with deep divergences. We identified four clades: i) one clade in coastal Georgia (referred as the Caucasus A lineage); ii) one distributed in Armenia and Crimea (Caucasus B lineage); iii) one is Israel (Israel lineage); and iv) one in SouthEastern Turkey, Iran and Armenia. We refer the latter as the *M. schaubi* lineage as it included the samples identified as *M. schaubi* from Armenia and Iran, which have a significantly larger body size. Surprisingly, the small form individuals from eastern Turkey also clustered within this lineage. The divergences among these four lineages varied between 3.4% (between the Israel lineage and the *M. schaubi* lineage) to 6.5% (between the Caucasus A lineage and the *M. schaubi* lineage).

### 3.2 Nuclear Intron Diversities

Among the analyzed nuclear markers, the intron 7 of the ROGDI gene was the least variable one (Figure 2 a; Table 1). It had very few variable sites and the haplotype and the nucleotide diversities were the lowest. The majority of the haplotypes were connected to a central one, which was present almost in all of the identified mtDNA clades, except the *M. escalerai* lineage. Despite to this relatively shallow divergence, some of the haplotypes were clade-specific; yet, the lineage sorting seemed to be far from being complete. The intron 5 of ABHD11 had relatively higher variability and the major lineages were mostly sorted. Though, one haplotype was shared among many clades, except the Western lineages (Figure 2 a). The intron 3 of ACOX2 and the intron 4 of COPS genes showed high variability and structuring. In both markers, the major mitochondrial lineages were almost sorted (Figure 2 a). In ACOX2, except for a few haplotypes, which were shared between the *M. nattereri* and the Central lineages, all the others were lineage specific. In COPS, the populations from the southern Anatolia and Israel shared a unique haplotype, which was connected to the *M. nattereri* lineage haplotypes.

**Table 1.** Polymorphism information about the nuclear introns; length after removing gaps (bp), samples size (N), number of variable sites (S), number of haplotypes (h), haplotype diversity (Hd), and nucleotide diversity (π).

| Gene | Length (bp) | N (this study) | S | h | Hd | $\pi$ |
| --- | --- | --- | --- | --- | --- | --- |
| ABHD11 intron 5 | 205 | 120 (86) | 20 | 21 | 0.834 | 0.015 |
| ACOX2 intron 3 | 372 | 118 (84) | 49 | 82 | 0.968 | 0.016 |
| COPS intron 4 | 599 | 90 (52) | 46 | 48 | 0.940 | 0.008 |
| ROGDI intron 7 | 340 | 115 () | 19 | 21 | 0.550 | 0.003 |

**FIGURE 1.**
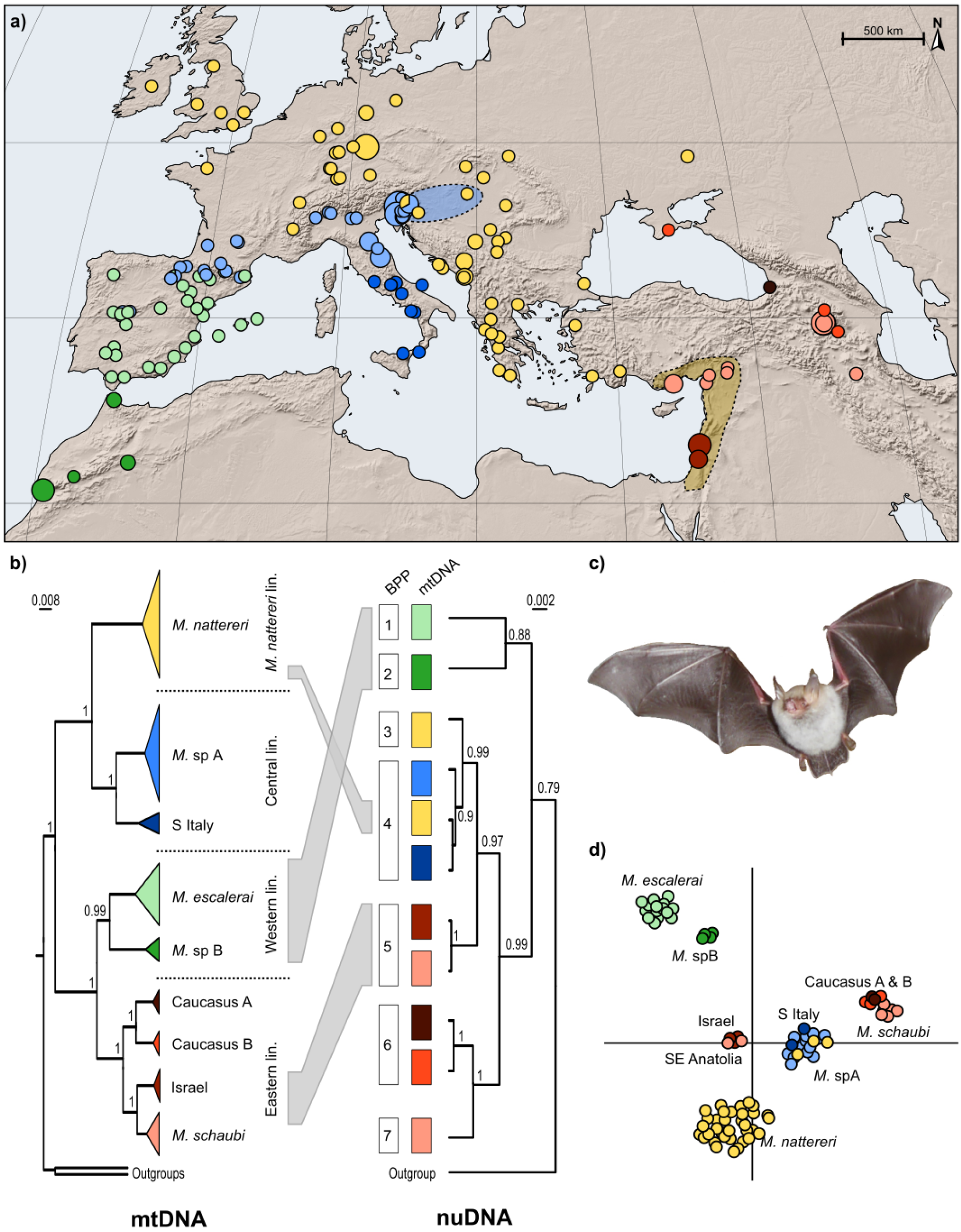
Distribution of the analyzed samples; shaded areas indicated areas of introgression (a). On the left, Bayesian tree of the partial ND1 gene; on the right, *Beast species tree based on nuclear introns, also showing the groups suggested by the BPP analysis. Grey connections indicate the discordant inferences between two genetic datasets: in nuDNA, i) the Western lineage located at the base of the tree; ii) few of the *M. nattereri* samples grouped with the Central lineage; and iii) the Israel lineage and the Anatolian populations of the the *M. schaubi* lineage clustered with the *M. nattereri* and the Central lineages (b). *M. nattereri* photo © Christian Dietz(c). The t-SNE visualization of the nuclear introns. Colored by the mtDNA lineage membership (d).

### 3.3 Population Genetic Structure

We used two different approaches to investigate the nuDNA structuring of the identified mitochondrial lineages. In PCA, we identified three major clusters which were mostly concordant with the mtDNA structure: i) the clades within the Western lineage clustered in one group; ii) the clades of the Eastern lineage clustered in another; and iii) the *M. nattereri* and the Central lineages formed another large group (Figure 2 b). The latter also included the populations from Israel and the southeastern Anatolia, which clustered within the Eastern lineage in mtDNA. We ran another PCA for this third group separately, which split them into three clusters: i) the *M. nattereri* lineage; ii) the Central lineage; and iii) the populations from Israel and the southeastern Anatolia (Figure 2 c). Here, some individuals from Slovenia and Hungary which belonged to the *M. nattereri* lineage in mtDNA clustered within the cluster of the Central lineage. The structure of these populations was more pronounced when the haplotype data was used (Figure S2.2). Here, the The t-SNE analysis revealed a very similar structure (Figure 1 d). Here, the *M. nattereri* and the Central lineages were more separated, and the latter contained again few individuals from the former. Similar to the PCA, *M. schaubi* lineage located very closely to the Caucasus A and Caucasus B lineages.

### 3.4 Species Tree Estimation and Delimitation

The *Beast and the BPP analyses estimated the same species tree (Figure 1 b). Its topology was different from the mtDNA tree. Firstly, in the nuDNA tree, the Western lineage located at the base of the tree. Secondly, similar to the PCA, the populations from Israel and the southeastern Anatolia clustered with the group of the *M. nattereri* and the Central lineages. Here, these populations were located at the basal split of this group, indicating that their separation was earlier than the split of the *M. nattereri* and the Central lineages. The BPP analysis grouped the identified lineages in to seven clusters (Figure 1 b). In addition to the previously suggested groups (*M. nattereri, M. escalerai, M.* sp A, *M.* sp B, and *M. schaubi*), here we identified two more new clusters. The first one is composed of the small form populations from the Caucasus and the Crimea. In nuDNA, this cluster was highly similar to *M. schaubi*, yet the latter has a substantially bigger body size (Horáček and Hanák 1984). The second new cluster is the group formed by the populations from Israel and the southeastern Anatolia.

## 4 Discussion

This study comprises all the recognized and the suggested species of the *Myotis nattereri* complex, covering almost their entire distribution range in the Western Palaearctic, except the *M*. sp. C from Corsica (Puechmaille et al. 2012). Three of these species, *M. nattereri, M. escalerai* and *M. schaubi*, can be identified by their distinct morphological traits (Horáček and Hanák 1984); and the other two, *M*. sp. A and *M*. sp. B, are proposed as distinct species based on their genetic differences (Salicini et al. 2011). Our dataset also includes samples from the populations of the eastern putative subspecies; *M. n. tschuliensis* from the Caucasus and *M. n. hoveli* from Israel (Horáček and Hanák 1984). In addition to previous studies (Puechmaille et al. 2012; Salicini et al. 2011, 2013), we identified other divergent lineages in the eastern ranges. Surprisingly, morphological classification did not always match the genetic relationships and the assignment of populations to the main genetic lineages differed between the mitochondrial and nuclear data sets.

### 4.1 Mitochondrial and nuclear discordances

In two areas, population assignments based on mitochondrial and nuclear markers did not fit with each other. Such conflicting patterns are observed when lineage sorting is incomplete or when introgression occurs (Toews and Brelsford 2012). We found the genetically discordant individuals always in the contact zones of distinct lineages. Such spatial patterns suggest that they resulted from introgression events rather than incomplete lineage sorting. Because, in the latter, discordant individuals would be expected to be randomly distributed rather than being geographically structured (Toews and Brelsford 2012).

**Table 2.**
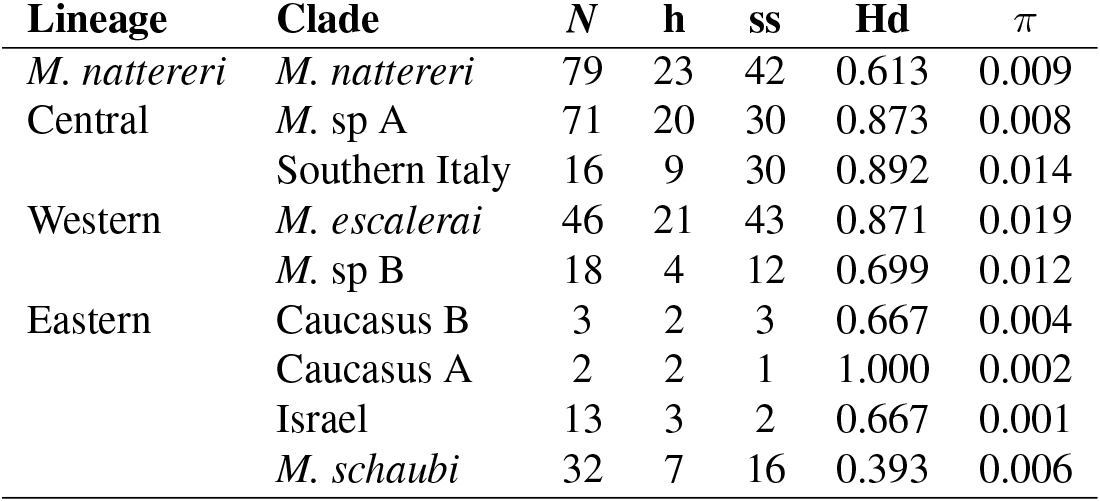
Summary statistics for the partial ND1 gene: number of samples (*N*), number of identified haplotypes (h), and segregating sites (ss), haplotype (Hd) and nucleotide (π) diversities.

**FIGURE 2.**
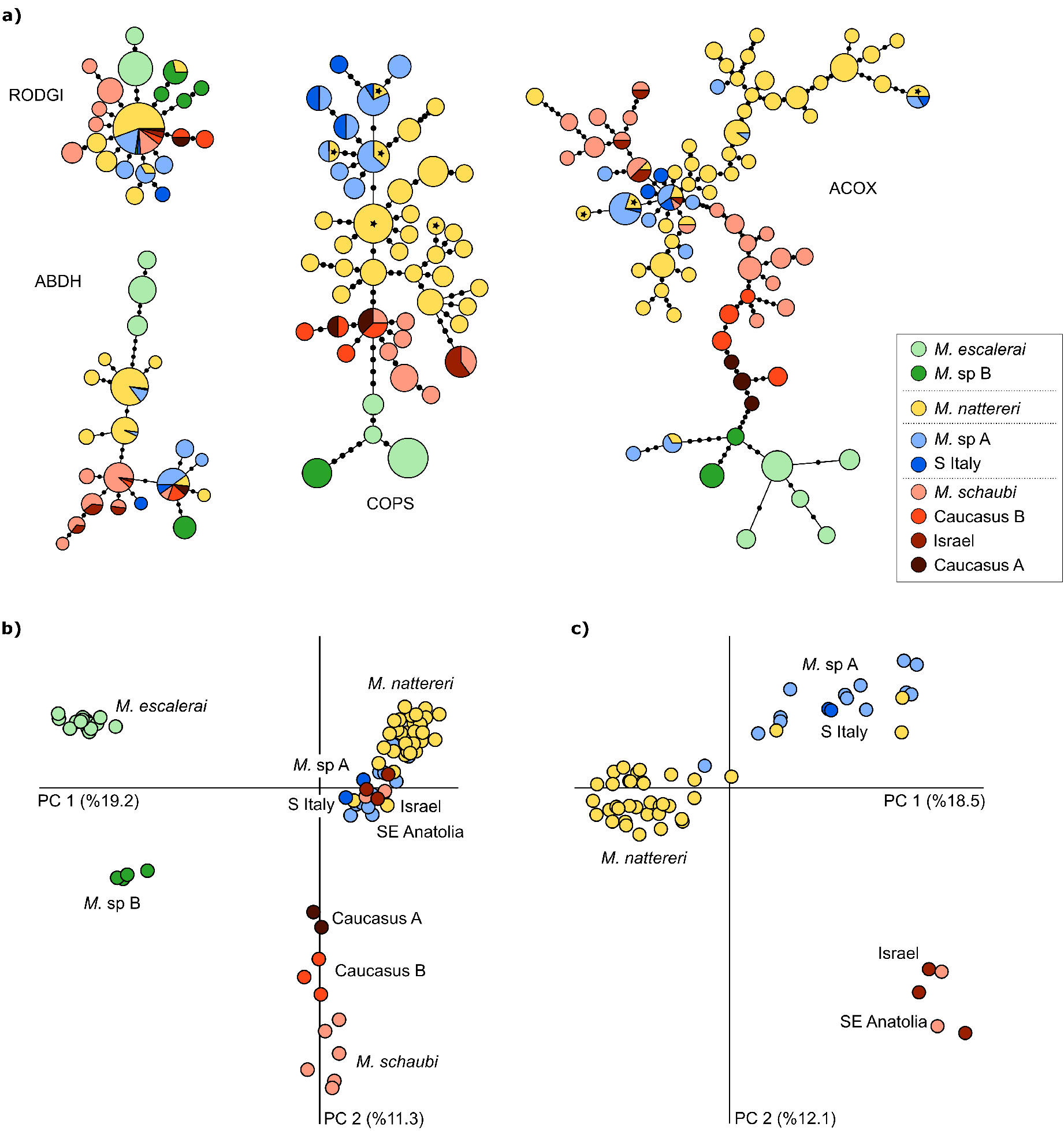
Population genetic structure based on nuclear introns. Haplotype networks; in ACOX2 and COPS genes, haplotypes marked with denotes the *M. nattereri* individuals which group within the Central lineage (a). PCA for the whole dataset (b). PCA for the populations of the *M. nattereri*, the *M*. sp A, the Southern Italy, the Israel, and the SE Anatolia of *M. schaubi* lineages (c). The percentage of variance explained by the principle components are shown in the parenthesis. Coloring is according to the mtDNA lineages as shown in the insert box.

### 4.2 Out of the Italian Peninsula

The most recent of the identified introgressions occurred between *M*. sp. A and *M. nattereri.* In mtDNA, these lineages have a contact zone in the northern coast of the Adriatic Sea, around Slovenia. We found that individuals in this area which belonged to *M*. *nattereri* in mtDNA clustered within the *M*. sp. A clade in nuclear analysis (Figure 1 b,d). This pattern can be explained by hybridization when *M*. sp. A expanded its range from the Apennines towards north and got into contact with *M*. *nattereri.* In the successive generations, back-crossing with the *M*. sp. A newcomers kept going on, shifting the nuclear genome of these populations towards *M*. sp. A, whereas, their mtDNA stayed as *M. nattereri*.

Because of the expansion history of these lineages, we consider that this introgression occurred relatively recently. The diversity and the distribution of the mtDNA haplotypes of *M. nattereri* suggest that its populations survived the last ice ages in the Balkans and the western coast of Anatolia. Populations from these areas have genetically diverse haplotypes and particularly three regions; western Anatolia, Greece, and the western Balkans, harbor high genetic diversity (Figure S2.1 a). On the other hand, in the western distribution range, the genetic diversity is low and individuals share one common haplotype or one of its closely related ones. This shallow diversity suggest that *M. nattereri* expanded its range relatively recently, probably just after the Last Glacial Maximum. Considering that the area of introgression is located in the middle of the expansion route of *M. nattereri*, we conclude that the hybridization occurred just after *M. nattereri* expanded its range to the west.

The level of current gene flow between these taxa remains unresolved. We did not identify any obvious hybrid individuals in our analysis. However, we had relatively few samples from the putative area of introgression. Further studies, focusing to this introgression area and its surroundings are needed to investigate the gene flow between these lineages. Considering that these two lineages are closely related, utilizing a genomic approach would be more effective to estimate the level of possible interbreeding.

### 4.3 Eastern Mediterranean Contact

In the southern coast of Anatolia and Israel, we identified other introgression events. In these areas, populations had two eastern mitochondrial lineages (Israel and *M. schaubi*), yet in nuclear DNA, they clustered into a single group, which was more related to the *M. nattereri* and Central lineages (Figure 1). These populations are likely to be descended from the hybrids which formed when *M. nattereri* expanded its range to the east and got into contact with the eastern lineages. Expanding *M. nattereri*, first, met with the *M. schaubi* lineage in southern Anatolia. They got the mtDNA of these local populations and later on, expanded further to the south, to Israel; once again, getting the mtDNA from the local populations.

The surprising aspect of both of these eastern introgressions, especially with the Israel lineage, was the lack of genetic clusters which could represent the ancestral populations. In nuclear DNA, we did not identify any distinct haplotypes which could represent the ancient local populations, from which the mitochondrial introgression occurred. Their nuclear genome might had been diminished and only a small portion of it might had been introgressed into the invading population of *M. nattereri.*

### 4.4 Body size evolution

Another interesting aspect of this introgression history is its possible relation to the body size evolution within this group. Despite their deep divergences, all the lineages are morphologically very similar, except *M. schaubi*, which is significantly larger. We consider that the larger body size is the derived phenotype. It is difficult to date when this increase happened. In mtDNA, the closest relatives of this large form are the Anatolian populations, which basically share the same mitochondrial clade. The next closest relative is the lineage from Israel. As the extant populations of these Eastern Mediterranean lineages are descended from the admixed populations with the small *M. nattereri* form, we cannot infer if their ancestral forms were small or large.

Considering the mitochondrial tree and the timing of the introgression events, the body size increase might have occurred in one of these three main periods: i) before the Israel lineage split; ii) after the Israel lineage split; and iii) after the introgression (Figure 3). In the first scenario, all the populations in the eastern Mediterranean would have a large body size and after their admixture with the *M. nattereri* lineage, they would have gotten smaller. Second scenario is similar, but only the *M. schaubi* lineage would have larger body size at the beginning. In both of these cases, the change in the body size might have been caused by an adaptive introgression.

In the final scenario, which we favor, we hypothesize that the body size increase happened only in the Caucasian populations of the *M. schaubi* lineage and relatively recently. Despite their deep divergence in the mtDNA, this group was closely related with the Caucasus A and B lineages in nuclear markers (Figure 1 b). This shallow divergence in nuclear DNA is an indication that the body size increase of *M. schaubi* occurred very recently and rather locally. Further studies focusing to the nuclear genomes of these taxa can provide details for the timing of body size evolution.

**Table 3.** Estimates of evolutionary divergence in partial ND1 gene over sequence pairs between groups shown in lower diagonal; their standard error estimates above the diagonal; and maximum divergences within groups in the diagonal.

| Lineage | Clade | 1 | 2 | 3 | 4 | 5 | 6 | 7 | 8 | 9 |
| --- | --- | --- | --- | --- | --- | --- | --- | --- | --- | --- |
| *M. nattereri* | 1) *M. nattereri* | **0.046** | 0.011 | 0.011 | 0.011 | 0.012 | 0.011 | 0.012 | 0.011 | 0.012 |
| Central | 2) *M. sp A* | 0.094 | **0.026** | 0.010 | 0.013 | 0.012 | 0.013 | 0.012 | 0.012 | 0.012 |
|  | 3) Southern Italy | 0.089 | 0.073 | **0.037** | 0.014 | 0.014 | 0.016 | 0.015 | 0.013 | 0.013 |
| Western | 4) *M. escaleraei* | 0.127 | 0.131 | 0.145 | **0.045** | 0.009 | 0.012 | 0.011 | 0.011 | 0.011 |
|  | 5) *M. sp B* | 0.119 | 0.110 | 0.139 | 0.082 | **0.022** | 0.012 | 0.011 | 0.010 | 0.010 |
| Eastern | 6) Caucasus A | 0.116 | 0.123 | 0.140 | 0.100 | 0.088 | **0.002** | 0.041 | 0.011 | 0.011 |
|  | 7) Caucasus B | 0.123 | 0.123 | 0.143 | 0.094 | 0.084 | 0.008 | **0.012** | 0.008 | 0.008 |
|  | 8) Israel | 0.115 | 0.117 | 0.132 | 0.091 | 0.086 | 0.058 | 0.043 | **0.003** | 0.007 |
|  | 9) *M. schaubi* | 0.121 | 0.121 | 0.134 | 0.101 | 0.090 | 0.065 | 0.050 | 0.034 | **0.024** |

**FIGURE 3.**
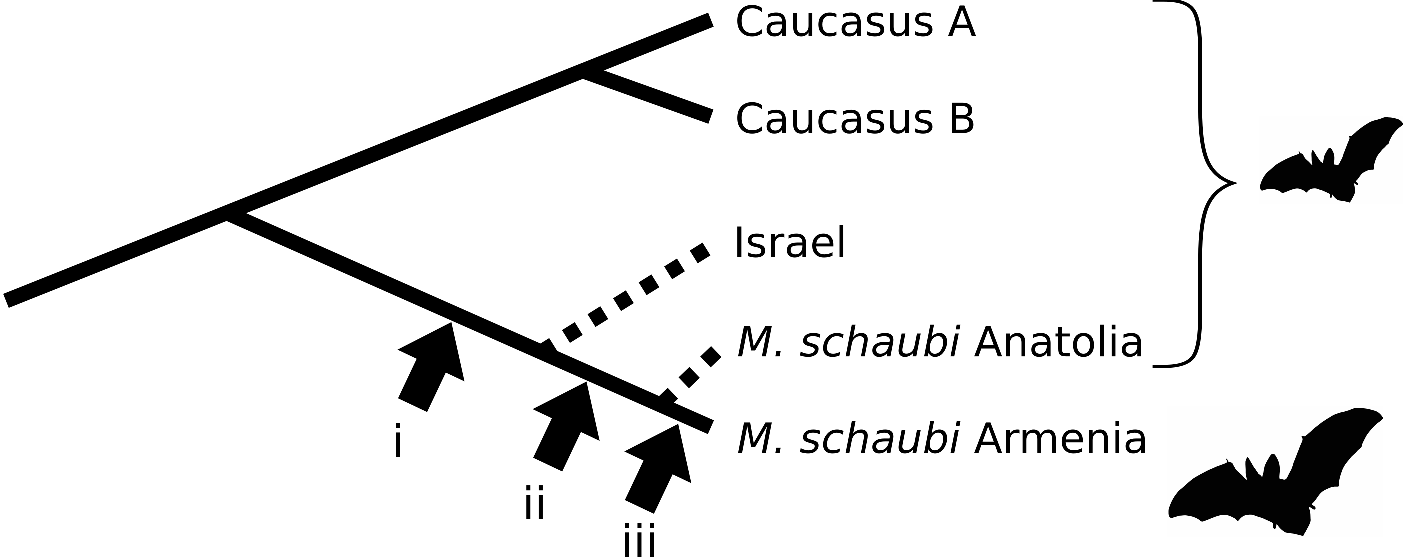
Possible scenarios for the body size evolution of the *M. schaubi*. The tree represents the eastern lineage branch from the mtDNA reconstruction in Figure 1, and the dashed lines indicates the lineages which were introgressed to *M. nattereri. Myotis schaubi* from Armenia has a larger body. Scenario: i) before the Israel lineage split; ii) after the Israel lineage split; and iii) after the introgression.

### 4.5 Conservation and taxonomic implications

Our study does not directly aim to resolve the taxonomic relations within this species complex. However, during our work, we were approached by various conservation practitioners, and were asked to comment on the taxonomic and accordingly, the conservation implications of the cryptic diversity within this group. It has been more than 10 years since the identification of some of these cryptic taxa (Ibáñez et al. 2006), and yet they still remain unnamed. In addition, here we present further cryptic taxa in the Caucasus, which could add to this unsettled nomenclature. Hence, we decided to infer the conservation and taxonomic implications of the genetic diversity within this species complex.

#### *M. nattereri helverseni* ssp. nov. Çoraman, Mayer and Dietz, 2018

We identified that there was a relatively recent mtDNA introgression from *M. nattereri* to *M*. sp A populations. This recent introgression suggest that these taxa are not reproductively isolated. Accordingly, we suggest that *M.* sp A represents a subspecies of *M. nattereri.* In reference to the late Otto von Helversen who was among the first to search systematically for cryptic diversity in European bats, we name this taxon as *M. nattereri helverseni* Çoraman, Mayer and Dietz 2018. As this group has a distinct gene pool, both for the nuclear and the mitochondrial DNA, it represents an evolutionarily significant unit (ESU), and should be considered in the conservation management plans.

#### *M. escalerai cabrerae* ssp. nov. Çoraman, Mayer and Dietz, 2018

The *M*. sp B lineage is clearly distinct, both in mitochondrial and nuclear markers. These distinct characters suggest that this taxon should be considered as an ESU, and accordingly, receive a separate protection plan. As this taxon is spatially separated from the rest of the lineages, it is not possible to infer if they are reproductively isolated. Accordingly, we suggest that this taxon should be considered as a subspecies of *M. escalerai*. We propose to name this taxon as *M. escalerai cabrerae* ssp. nov. Çoraman, Mayer and Dietz 2018 in reference to Angel Cabrera, who described *M. escalerai* from Spain (Ibáñez and Fernindez 2008).

#### *M. tschuliensis* sp. nov. Kuzyakin, 1935

Our findings suggest that the Caucasus hosts further cryptic diversity. Although populations of the ‘small form’ in this region morphologically closely resemble the European *M. nattereri*, in terms of their mitochondrial and nuclear genes, they are more closely related to the ‘large form’ *M. schaubi.* These two forms occur in close proximity in Armenia. They differ substantially in their body size. In Armenia, we caught both species only 90 km apart from each other. Despite the close geographic proximity we did not catch animals of intermediate body size. The lack of mitochondrial introgression, coupled with the lack of intermediate phenotypes argue for the existence of effective hybridization barriers and thus distinct biological species. Accordingly, we suggest assigning these Caucasian ‘small form’ populations as *M. tschuliensis* Kuzyakin, 1935, following the formerly proposed sub-specific nomination (Benda et al. 2006, 2012, 2011; Horáček and Hanák 1984).

#### *M. hoveli* sp. nov. Harrison, 1964

Populations from Israel and the southeastern Anatolia formed a distinct cluster in nuDNA. This group had its mtDNA from the eastern lineages, yet clustered with the *M. nattereri* and the central lineages in nuDNA. As we did not identify any admixture with the neighboring lineages, we consider that these populations represent a distinct species and name them tentatively as *M. hoveli* Harrison, 1964, referring to the formerly proposed sub-specific nomination of the populations from Israel (Horáček and Hanák 1984).

#### *M. araxenus* sp. nov. Dahl, 1974

As the large and the small forms diverged relatively recently, we also suggest to change the name of *M. schaubi.* The populations from the Caucasus which had larger body size were named as *M. schaubi* due to their morphological similarity to a Pliocene fossil found in Hungary (Horáček and Hanák 1984). Our findings suggest that this larger form in the lower Caucasus and Iran probably evolved in this area and rather recently. In that case, *M. schaubi* described in Hungary probably represents a distinct form; likely to be an ancestral form of the whole group. Therefore, we propose to rename the large form populations as *M. araxenus*, following their prior subspecific description, *M. nattereri araxenus* Dahl, 1947 (Horáček and Hanák 1984).

**FIGURE 4.**
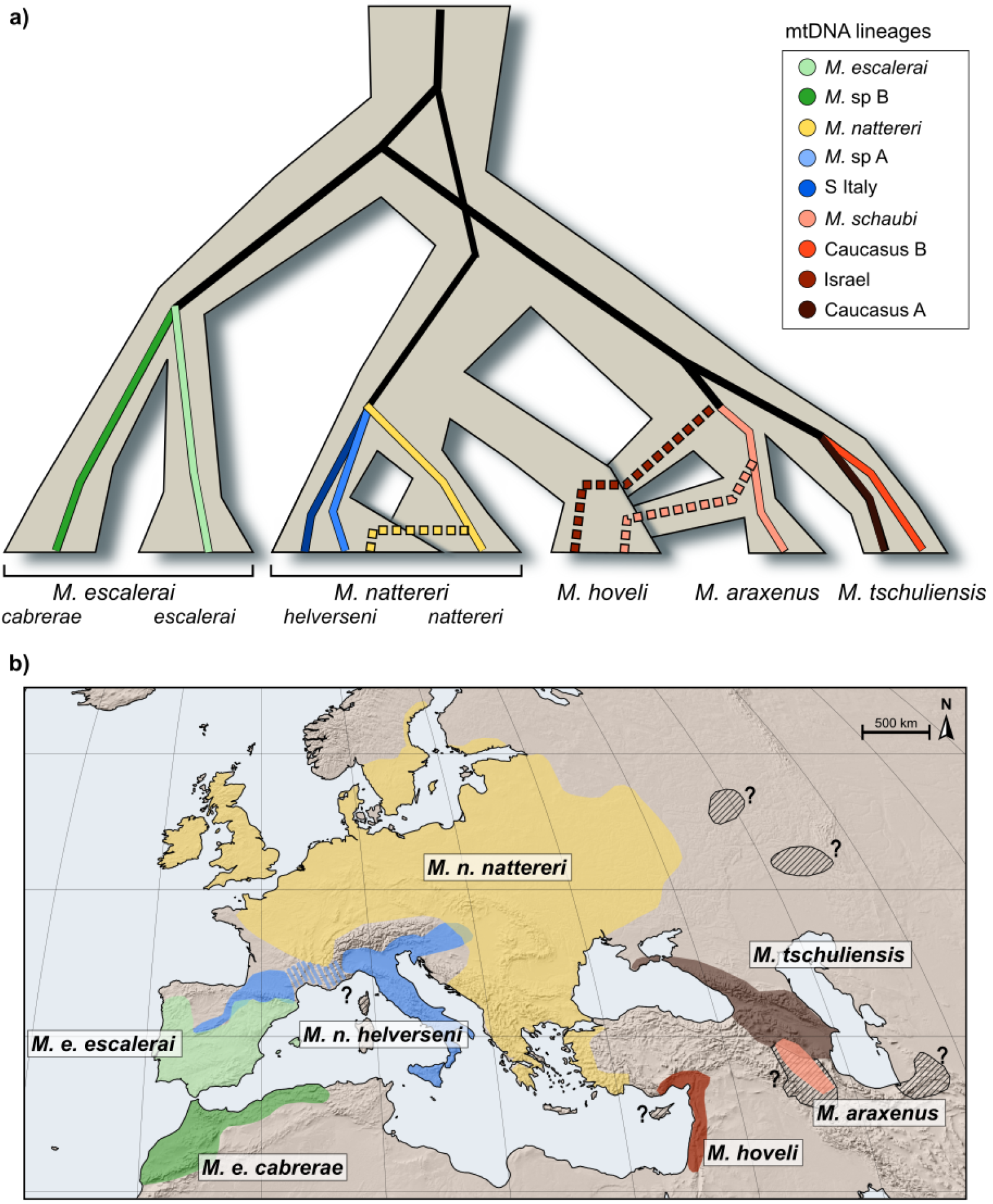
Graphical representation of the reticulate evolutionary history of *M. nattereri* sensu lato. Grey tree shows the inferred species tree and the lines indicate mitochondrial lineages; dashed ones show introgression events. Note that, in the case of *M. hoveli*, introgressions probably occurred in reverse order or relatively at the same time. Because of the graphical clarity, we present as it is (a). Distribution of taxa based on the suggested revisions. Dashed areas indicate unsampled distribution ranges (b).

### 4.6 Reticulating histories and extinct lineages

Our study suggests that admixture events played a major role in the evolutionary history of the *M. nattereri* complex (Figure 4 a). Nowadays, there is increasing evidence showing that species within various taxonomic groups had gene flow after their divergences. For instance, big cats (Figueiró et al. 2017), elephants (Palkopoulou et al. 2018), and humans (Prüfer et al. 2014) have reticulating phylogenetic trees rather than having simple bifurcating relationships. The patterns of mtDNA introgressions among the lineages of the *M. nattereri* complex suggest a similar reticulating evolutionary history. Lineages within this group contracted and expanded their ranges multiple times and frequently got into contact in different areas. In some of these contacts, populations hybridized and exchanged genetic material.

An interesting finding of our study is that one of the identified mtDNA lineages was introgressed from a taxon which probably got extinct. This taxon might have been driven to extinction because of the competition with the invading population, to whom the mtDNA introgressed, or because of other factors, such as changing environmental conditions. This scenario is similar to the evolutionary history of humans (Prüfer et al. 2014). When humans expanded out of Africa, they got into contact and admixed with Neanderthals and Denisovans. Modern non-African humans inherit a small fraction of their genome from them. Both of these taxa are extinct, yet their genes survive in our genomes. To our knowledge, the Israel mitochondrial lineage in the *M. nattereri* complex is one of the very few examples where an extinct taxon was identified based on only its surviving DNA in another species (see Xu et al. (2017), for an example of extinct archaic hominin).

The ‘disappearance’ of these ancestral populations might be also explained by fusion of the lineages. In such a scenario, the local and the invading populations fuse into a single lineage, forming a new taxon, which has a mosaic genome of its parental forms (Jacobsen and Omland 2011). Lineage fusion or hybrid speciation, when the fused lineages form a distinct species, have been documented within various species groups (Garrick et al. 2014; Kearns et al. 2018; Kleindorfer et al. 2014; Payseur and Rieseberg 2016; Vonlanthen et al. 2012). Further studies, focusing on the genomic landscape of the *M. hoveli* populations can provide insights about their reticulation history.

It is likely that the admixture events which led to the observed patterns of introgressions occurred when climate was changing (see Cahill et al. (2018) for an example). During such periods, distribution of habitats and accordingly the associated taxa reshape, often bringing together formerly isolated species (Currat et al. 2008; Hewitt 2000). Hybridization occurring at such periods can provide evolutionary novelty into gene pools of these expanding species, which in turn can affect their fitness (Cahill et al. 2018). Nowadays, complex reticulate histories are emerging all across the tree of life (Cahill et al. 2018; Palkopoulou et al. 2018; Prüfer et al. 2014). We think that understanding the biogeographical history of such species complexes will provide crucial insight about the underlying mechanisms of reticulated trees. Here, we show that dense sampling and analysis of nuclear markers can reveal complex evolutionary histories with multiple contact zones and various hybridization events.

## ACKNOWLEDGEMENTS

We are grateful to Isabelle Waurick; she did the DNA extractions and performed sequencing. We would like to thank Benjamin Allegrini, Yalin Emek Çelik, Isabel Dietz, Fulgencio Lison Gil, Stefan Greif, Tamás Görföl, Matej Hočcevar, Otto von Hel-versen, Richard Hoffman, Maria Jerabek, Darija Josić, Andreas Kiefer, Jana Mlakar, Bernd Ohlendorf, Eleni Papadatou, George Popov, Boyan Petrov, Alenka Petrinjak, Monika Podgorelec, Guido Reiter, Simon Ripperger, Bernd-Ulrich Rudolph, Konrad Sachanowicz, Marjetka Šemrl, Aleš Tomažič, Gudrun Wibbelt, Anton Vlaschenko, Andreas Zahn, and various other researchers for helping during the field works and/or getting access to tissue samples. MZ was funded by the Slovenian research agency through the Research program P1-0184.

## Bibliography

Benda, P., Andreas, M., Kock, D., Lucan, R. K., Munclinger, P., Nova, P., Obuch, J., Ochman, K., Reiter, A., and Uhrin, M. (2006). Bats (Mammalia: Chiroptera) of the Eastern Mediterranean. Part 4. Bat fauna of Syria: distribution, systematics, ecology. Acta Societatis Zoologicae Bohemicae, 70(1):1–329.

Benda, P., Faizolâhi, K., Andreas, M., Obuch, J., Reiter, A., Ševčík, M., Uhrin, M., Vallo, P., and Ashrafi, S. (2012). Bats (Mammalia: Chiroptera) of the Eastern Mediterranean and Middle East. Part 10. Bat fauna of Iran. Acta Societatis Zoologicae Bohemicae, 76:163–582.

Benda, P., Hanák, V., and ČervenÝ, J. (2011). Bats (Mammalia: Chiroptera) of the Eastern Mediterranean and Middle East. Part 9. Bats from Transcaucasia and West Turkestan in collection of the National Museum, Prague. Acta Societatis Zoologicae Bohemicae, 75:159–222.

Cahill, J. A., Heintzman, P D., Harris, K., Teasdale, M. D., Kapp, J., Soares, A. E. R., Stirling, I., Bradley, D., Edwards, C. J., Graim, K., Kisleika, A. A., Malev, A. V., Monaghan, N., Green, R. E., Shapiro, B., and Su, B. (2018). Genomic evidence of widespread admixture from polar bears into brown bears during the last ice age. Molecular Biology and Evolution, 35(5):1120–1129.

Currat, M., Ruedi, M., Petit, R. J., Excoffier, L., and Wolf, J. (2008). The hidden side of invasions: Massive introgression by local genes. Evolution, 62(8):1908–1920.

Çoraman, E., Furman, A., Karatas, A., and Bilgin, R. (2013). Phylogeographic analysis of Anatolian bats highlights the importance of the region for preserving the Chiropteran mitochondrial genetic diversity in the Western Palaearctic. Conservation Genetics, 14(6):1205–1216.

Darriba, D., Taboada, G. L., Doallo, R., and Posada, D. (2012). jModelTest 2: more models, new heuristics and parallel computing. Nature Methods, 9(8):772–772.

Dietz, C., Gazaryan, A., Papov, G., Dundarova, H., and Mayer, F. (2016). Myotis hajastanicus is a local vicariant of a widespread species rather than a critically endangered endemic of the Sevan lake basin (Armenia). Mammalian Biology-Zeitschrift für Säugetierkunde, 81(5):518–522.

Drummond, A. J., Suchard, M. A., Xie, D., and Rambaut, A. (2012). Bayesian phylogenetics with BEAUti and the BEAST 1.7. Molecular Biology and Evolution, 29(8):1969–1973.

Figueiró, H. V., Li, G., Trindade, F. J., Assis, J., Pais, F., Fernandes, G., Santos, S. H. D., Hughes, G. M., Komissarov, A., Antunes, A., Trinca, C. S., Rodrigues, M. R., Linderoth, T, Bi, K., Silveira, L., Azevedo, F. C. C., Kantek, D., Ramalho, E., Brassaloti, R. A., Villela, P. M. S., Nunes, A. L. V., Teixeira, R. H. F., Morato, R. G., Loska, D., Saragüeta, P, Gabaldón, T, Teeling, E. C., O’Brien, S. J., Nielsen, R., Coutinho, L. L., Oliveira, G., Murphy, W. J., and Eizirik, E. (2017). Genome-wide signatures of complex introgression and adaptive evolution in the big cats. Science Advances, 3(7):e1700299.

Garrick, R. C., Benavides, E., Russello, M. A., Hyseni, C., Edwards, D. L., Gibbs, J. P, Tapia, W., Ciofi, C., and Caccone, A. (2014). Lineage fusion in Galápagos giant tortoises. Molecular Ecology, 23(21):5276–5290.

Heled, J. and Drummond, A. J. (2009). Bayesian inference of species trees from multilocus data. Molecular Biology and Evolution, 27(3):570–580.

Hewitt, G. (2000). The genetic legacy of the Quaternary ice ages. Nature, 405(6789):907–913.

Horáček, I. and Hanák, V. (1984). Comments on the systematics and phylogeny of myotis nattereri (Kuhl, 1818). Myotis, 21(22):20–29.

Ibáñez, C. and Fernández, R. (2008). Catálogo de murciélagos de las colecciones del Museo Nacional de Ciencias Naturales. Monografías Museo Nacional de Ciencias Naturales, 2:1–54.

Ibáñez, C., García-Mudarra, J. L., Ruedi, M., Stadelmann, B., and Juste, J. (2006). The Iberian contribution to cryptic diversity in European bats. Acta Chiropterologica, 8(2):277–297.

Jacobsen, F. and Omland, K. E. (2011). Increasing evidence of the role of gene flow in animal evolution: hybrid speciation in the yellow-rumped warbler complex. Molecular Ecology, 20(11):2236–2239.

Jombart, T. (2008). adegenet: a R package for the multivariate analysis of genetic markers. Bioinformatics, 24(11):1403–1405.

Kearns, A. M., Restani, M., Szabo, I., Schrøder-Nielsen, A., Kim, J. A., Richardson, H. M., Marzluff, J. M., Fleischer, R. C., Johnsen, A., and Omland, K. E. (2018). Genomic evidence of speciation reversal in ravens. Nature Communications, 9(1):906.

Kleindorfer, S., O’Connor, J. A., Dudaniec, R. Y, Myers, S. A., Robertson, J., and Sulloway, F. J. (2014). Species collapse via hybridization in Darwin’s tree finches. The American Naturalist, 183(3):325–341.

Kumar, S., Stecher, G., and Tamura, K. (2016). MEGA7: Molecular Evolutionary Genetics Analysis version 7.0 for bigger datasets. Molecular Biology and Evolution, 33(7):1870–1874.

Li, G., Davis, B. W., Eizirik, E., and Murphy, W. J. (2016). Phylogenomic evidence for ancient hybridization in the genomes of living cats (Felidae). Genome Research, 26(1):1–11.

Li, W., Cerise, J. E., Yang, Y, and Han, H. (2017). Application of t-SNE to human genetic data. Journal of Bioinformatics and Computational Biology, 15(04):1750017.

Librado, P. and Rozas, J. (2009). DnaSP v5: a software for comprehensive analysis of DNA polymorphism data. Bioinformatics, 25(11):1451–1452.

van der Maaten, L. and Hinton, G. (2008). Visualizing data using t-SNE. Journal of Machine Learning Research, 9(Nov):2579–2605.

Palkopoulou, E., Lipson, M., Mallick, S., Nielsen, S., Rohland, N., Baleka, S., Karpinski, E., Ivancevic, A. M., To, T.-H., Kortschak, R. D., Raison, J. M., Qu, Z., Chin, T.-J., Alt, K. W., Claesson, S., Dalén, L., MacPhee, R. D. E., Meller, H., Roca, A. L., Ryder, O. A., Heiman, D., Young, S., Breen, M., Williams, C., Aken, B. L., Ruffier, M., Karlsson, E., Johnson, J., Palma, F. D., Alfoldi, J., Adelson, D. L., Mailund, T., Munch, K., Lindblad-Toh, K., Hofreiter, M., Poinar, H., and Reich, D. (2018). A comprehensive genomic history of extinct and living elephants. Proceedings of the National Academy of Sciences. https://doi.org/10.1073/pnas.1720554115

Paradis, E. (2010). pegas: an R package for population genetics with an integrated-modular approach. Bioinformatics, 26(3):419–420.

Payseur, B. A. and Rieseberg, L. H. (2016). A genomic perspective on hybridization and speciation. Molecular Ecology, 25(11):2337–2360.

Petit, R. J. and Excoffier, L. (2009). Gene flow and species delimitation. Trends in Ecology & Evolution, 24(7):386–393.

Prüfer, K., Racimo, F., Patterson, N., Jay, F., Sankararaman, S., Sawyer, S., Heinze, A., Renaud, G., Sudmant, P. H., de Filippo, C., Li, H., Mallick, S., Dannemann, M., Fu, Q., Kircher, M., Kuhlwilm, M., Lachmann, M., Meyer, M., Ongyerth, M., Siebauer, M., Theunert, C., Tandon, A., Moorjani, P., Pickrell, J., Mullikin, J. C., Vohr, S. H., Green, R. E., Hellmann, I., Johnson, P L. F., Blanche, H., Cann, H., Kitzman, J. O., Shendure, J., Eichler, E. E., Lein, E. S., Bakken, T. E., Golovanova, L. V., Doronichev, V. B., Shunkov, M. V, Derevianko, A. P., Viola, B., Slatkin, M., Reich, D., Kelso, J., and Pääbo, S. (2014). The complete genome sequence of a Neanderthal from the Altai Mountains. Nature, 505(7481):43–49.

Puechmaille, S. J., Allegrini, B., Boston, E. S., Dubourg-Savage, M.-J., Evin, A., Knochel, A., Le Bris, Y, Lecoq, V., Lemaire, M., and Rist, D. (2012). Genetic analyses reveal further cryptic lineages within the Myotis nattereri species complex. Mammalian Biology-Zeitschrift für Säugetierkunde, 77(3):224–228.

Rannala, B. and Yang, Z. (2003). Bayes estimation of species divergence times and ancestral population sizes using DNA sequences from multiple loci. Genetics, 164(4):1645–1656.

Rheindt, F. E. and Edwards, S. V. (2011). Genetic introgression: An integral but neglected component of speciation in birds. The Auk, 128(4):620–632.

Ruedi, M. and Mayer, F. (2001). Molecular systematics of bats of the genus Myotis (Vespertilionidae) suggests deterministic ecomorphological convergences. Molecular Phylogenetics and Evolution, 21(3):436–448.

Salicini, I., Ibáñez, C., and Juste, J. (2011). Multilocus phylogeny and species delimitation within the Natterer’s bat species complex in the Western Palearctic. Molecular Phylogenetics and Evolution, 61(3):888–898.

Salicini, I., Ibáñez, C., and Juste, J. (2013). Deep differentiation between and within Mediterranean glacial refugia in a flying mammal, the Myotis nattereri bat complex. Journal of Biogeography, 40(6):1182–1193.

Salicini, I., Ibáñez, C., and Juste, J. (2018). Corrigendum to “Multilocus phylogeny and species delimitation within the Natterer’s bat species complex in the Western Palearctic” [Molecular Phylogenetics and Evolution 61 (2011) 888-898]. Molecular Phylogenetics and Evolution, 120:391–392.

Takahata, N., Satta, Y, and Klein, J. (1995). Divergence time and population size in the lineage leading to modern humans. Theoretical Population Biology, 48(2):198–221.

Toews, D. P. L. and Brelsford, A. (2012). The biogeography of mitochondrial and nuclear discordance in animals. Molecular Ecology, 21(16):3907–3930.

Vonlanthen, P., Bittner, D., Hudson, A. G., Young, K. A., Müller, R., Lundsgaard-Hansen, B., Roy, D., Di Piazza, S., Largiader, C. R., and Seehausen, O. (2012). Eutrophication causes speciation reversal in whitefish adaptive radiations. Nature, 482(7385):357–362.

Wallis, G. P., Cameron-Christie, S. R., Kennedy, H. L., Palmer, G., Sanders, T. R., and Winter, D. J. (2017). Interspecific hybridization causes long-term phylogenetic discordance between nuclear and mitochondrial genomes in freshwater fishes. Molecular Ecology, 26(12):3116–3127.

Xu, D., Pavlidis, P, Taskent, R. O., Alachiotis, N., Flanagan, C., DeGiorgio, M., Blekhman, R., Ruhl, S., and Gokcumen, O. (2017). Archaic Hominin introgression in Africa contributes to functional salivary MUC7 genetic variation. Molecular Biology and Evolution, 34(10):2704–2715.

Yang, Z. (2002). Likelihood and Bayes estimation of ancestral population sizes in hominoids using data from multiple loci. Genetics, 162(4):1811–1823.

Yang, Z. (2015). The BPP program for species tree estimation and species delimitation. Current Zoology, 61(5):854–865.

